# Gkmexplain: Fast and Accurate Interpretation of Nonlinear Gapped *k*-mer SVMs Using Integrated Gradients

**DOI:** 10.1101/457606

**Authors:** Avanti Shrikumar, Eva Prakash, Anshul Kundaje

**Affiliations:** Stanford University; BASIS Silicon Valley

## Abstract

Support Vector Machines with gapped *k*-mer kernels (gkm-SVMs) have been used to learn predictive models of regulatory DNA sequence. However, interpreting predictive sequence patterns learned by gkm-SVMs can be challenging. Existing interpretation methods such as deltaSVM, in-silico mutagenesis (ISM), or SHAP either do not scale well or make limiting assumptions about the model that can produce misleading results when the gkm kernel is combined with nonlinear kernels. Here, we propose gkmexplain: a novel approach inspired by the method of Integrated Gradients for interpreting gkm-SVM models. Using simulated regulatory DNA sequences, we show that gkmexplain identifies predictive patterns with high accuracy while avoiding pitfalls of deltaSVM and ISM and being orders of magnitude more computationally efficient than SHAP. We use a novel motif discovery method called TF-MoDISco to recover consolidated TF motifs from gkm-SVM models of *in vivo* TF binding by aggregating predictive patterns identified by gkmexplain. Finally, we find that mutation impact scores derived through gkmexplain using gkm-SVM models of chromatin accessibility in lymphoblastoid cell-lines consistently outperform deltaSVM and ISM at identifying regulatory genetic variants (dsQTLs). Code and example notebooks replicating the workflow are available at https://github.com/kundajelab/gkmexplain. Explanatory videos available at http://bit.ly/gkmexplainvids.

## 1 Introduction

Deciphering the combinatorial regulatory DNA sequence patterns that determine transcription factor (TF) binding and chromatin state is critical to understand gene regulation and interpret the molecular impact of regulatory genetic variation. High-throughput *in vivo* and *in vitro* functional genomics experiments provide large datasets to train predictive models using machine learning approaches that can learn the relationship between regulatory DNA sequences and their associated molecular phenotypes. Support Vector Machines (SVMs) are a popular choice in machine learning prediction tasks because they are stable to train and, when used with an appropriate kernel, can model complex input-output relationships. For classification tasks, SVMs learn an optimal linear separating hyperplane in a high-dimensional space determined by a kernel that measures the similarity between all pairs of examples. The gapped *k*-mer (gkm) string kernel [11, 6] was developed to enable training SVMs on string inputs such as DNA or protein sequences. The gkm kernel computes a similarity between pairs of sequences based on biologically motivation notion of shared approximate occurrences of short subsequences allowing for gaps and mismatches. The gkm kernel can be further combined with other nonlinear kernels such as radial basis function (RBF) kernel to capture complex nonlinear relationships between the input string features. Gapped *k*-mer SVMs and their extensions have been successfully applied to several prediction tasks in regulatory genomics such as TF binding and chromatin accessibility prediction. Interpretation of these trained SVMs is important to decipher the patterns such as TF binding motifs present in the any DNA sequences that are predictive of its associated molecular label. Moreover, these models can be used to predict the impacts of genetic variants in regulatory DNA sequences.

Unfortunately, interpretation of gkm-SVMs can be challenging. DeltaSVM [9], a popular tool developed by the authors of gkm-SVM to estimate the effects of variants, can produce unsatisfactory results when used with nonlinear versions of the gkm kernel such as the gkmrbf kernel [8]. In-silico mutagenesis (ISM) [4], an interpretation approach that consists of performing individual perturbations and observing the impact on the output, can be computationally inefficient and can fail to reveal the presence of motifs when nonlinear saturation effects are present [13]. SHAP [12], a method based on estimating Shapely values, can address the nonlinearity issues faced by deltaSVM and ISM but is even less computationally efficient than ISM. As such, there is a need for fast interpretation of gapped *k*-mer Support Vector Machines that works reliably in the presence of nonlinear effects.

Here we present gkmexplain, a computationally efficient method for explaining the predictions of both linear and nonlinear variants of gapped *k*-mer SVMs. Specifically, gkmexplain is a feature attribution method that efficiently computes the predictive contribution or importance of every nucleotide in an input DNA sequence to its associated output label through the lens of a gkmSVM model. gkmexplaim works by decomposing the output of the gapped *k*-mer string kernel into the contributions of matching positions within the *k*-mers, and can be interpreted as a variation of the Integrated Gradients, a feature attribution method developed to interpret deep learning models [15]. On simulated regulatory DNA sequences, where the ground truth predictive motifs are known, we show that gkmexplain outperforms deltaSVM and ISM while being multiple orders of magnitude more computationally efficient than SHAP. We use a novel motif discovery method called TF-MoDISco [14] to recover consolidated TF motifs from gkm-SVM models of *in vivo* TF binding by aggregating predictive patterns identified by gkmexplain. Finally, using non-linear gkmSVM models of chromatin accessible regulatory DNA sequences, we show that mutation impact scores derived through gkmexplain outperform deltaSVM and ISM at identifying DNAse-I hypersensitive QTLs (dsQTLs) in lymphoblastoid cell-lines.

## 2 Background

### 2.1 Gapped *k*-mers

A full *k*-mer refers to a letter subsequence of length *k* - for example, AAGT is a full 4-mer. By contrast, a gapped *k*-mer refers to a subsequence containing *k* letters and some number of gaps - for example, A*AG*T is a gapped 4-mer containing 2 gaps (* is used to denote a gap). In the gkm-SVM implementation, *l* denotes the full length of the subsequences considered (including gaps), and *k* denotes the number of non-gap positions - for example, the *l*-mer ACG (where *l* = 3) contains the gapped *k*-mers AC*, A*G and *CG (where *k* = 2).

### 2.2 Support Vector Machines and Gapped *k*-mer Kernels

Let **x** be an input to a Support Vector Machine (SVM), *m* be the total number of support vectors, *Z^i^* be the *i*th support vector, *y^i^* be the label (+1 or -1) associated with the *i*th support vector, *α_i_* be the weight associated with the *i*th support vector, *b* be a constant bias term, and *K* be a *kernel function* that is used to compute a similarity score between *Z^i^* and **x**. The SVM produces an output of the form:

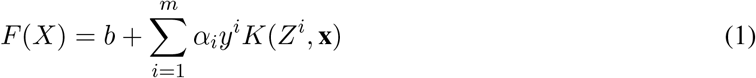

Kernel functions can be thought of as implicitly computing dot products between their inputs mapped to vectors in some feature space. For example, the *gapped k-mer* kernel implicitly maps its inputs to feature vectors representing the normalized counts of gapped *k*-mers. Thus, it has the form:

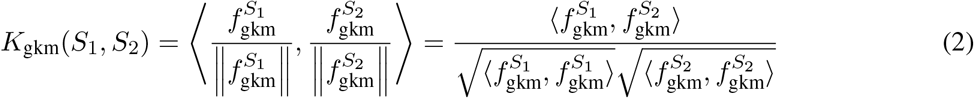

Where 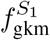 and 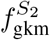 are feature vectors consisting of the counts of gapped *k*-mers in sequences *S*_1_ and *S*_2_ respectively. Because the feature space corresponding to gapped *k*-mer counts can be quite large, a more computationally efficient formulation [11, 6] of the gapped *k*-mer kernel performs a sum only over the full *l*-mers that are actually present in the sequences. Let 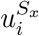 represent the identity of the *l*-mer at position *i* in sequence *S_x_*, and let 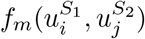 be a function that returns the number of mismatches between the *l*-mers 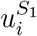 and 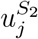. We can write 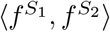 as:

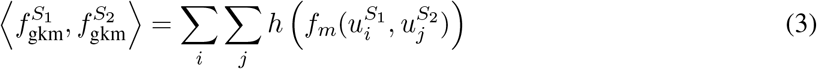

Where the indexes *i* and *j* sum over all *l*-mers in *S*_1_ and *S*_2_ respectively, and *h*(*m*) is a function that returns the contribution of an *l*-mer pair with *m* mismatches to the dot product 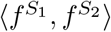. For example, if 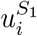and 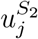 are a pair of *l*-mers with *m* mismatches between them, then the number of gapped *k*-mers they share is 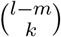. Thus, in the case of the traditional gapped *k*-mer kernel, 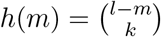 Note that Gandhi et al. [6] proposed variants of the gapped *k*-mer kernel such as the *truncated gkm-filter*. These variants differ in the function *h*(*m*), but otherwise have an identical formulation.

### 2.3 Extensions of the gkm kernel

We described the extensions to the gkm kernel proposed by Lee in [8].

#### 2.3.1 The wgkm kernel

In the weighted gkm (wgkm) kernel, *l*-mers are given a weight according to the position at which they occur. A dot product in the feature space of weighted gapped *k*-mers can thus be expressed as:

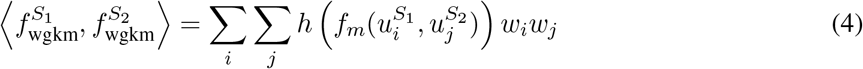

Where *w_i_* and *w_j_* are the weights associated with *l*-mers at position *i* and *j* respectively. The weighted gkm kernel then becomes:

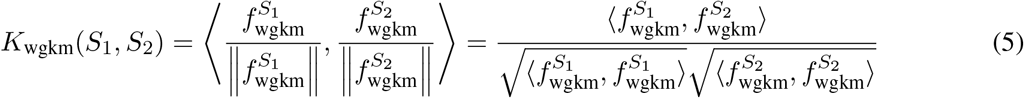

Note that the gkm kernel is a special case of the wgkm kernel with *w_i_* = 1 for all *i*.

#### 2.3.2 The gkmrbf kernel

The RBF kernel is useful for modeling complex nonlinear interactions between input features, and is defined as 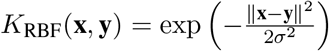 (where **x** and **y** are vectors). Recall that the gkm kernel can be thought of as mapping the input sequences to a feature space of normalized gapped *k*-mer counts. The gkmrbf kernel maps sequences to the same feature space as the gkm kernel, but then applies the RBF kernel rather than the dot product. For efficient computation, we leverage the following equality:

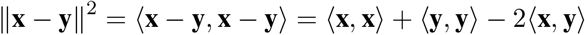

As gkm feature vectors are always unit normalized, 〈**x**, **x**〉 = 〈**y**, **y**〉 = 1. If we let 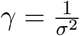, we get:

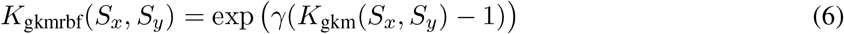

#### 2.3.3 The wgkmrbf kernelThe

Analogous to **Eqn. 6**, the wgkmrbf kernel is a combination of the wgkm kernel and the RBF kernel:

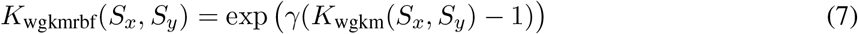

## 3 Previous Work

DeltaSVM [9] is a popular tool developed by the authors of gkm-SVM to estimate the in-silico effects of genetic variants. DeltaSVM defines a score for each *l*-mer as the output of the gkm-SVM when the gkm-SVM is supplied only that *l*-mer as input. It then estimates the impact of a mutation as the total change in the scores of all *l*-mers overlapping the mutation. Although computationally efficient, this formulation does not consider interactions between *l*-mers that can be learned by the gkmrbf or wgkmrbf kernels (**Sec. 2.3**). It also does not consider the influence of positional weights that are present in the wgkm or wgkmrbf kernels. Consistent with this, Lee [8] observed that deltaSVM used in conjunction with the gkmrbf, wgkm or wgkmrbf kernels did not produce improvements in variant scoring relative to the gkm kernel, even though predictive models trained with the former kernels performed better.

An alternative to deltaSVM for estimating the impact of individual mutations is to directly compute the change in the output of the SVM when the mutation is introduced into the input sequence. This approach is called in-silico mutagenesis (ISM) [4], and it has been successfully applied to interpret complex machine learning models [17]. However, individually computing the impact of all possible mutations in an input sequence can become computationally inefficient. Further, ISM can fail to reveal the presence of motifs in a sequence when nonlinear saturating effects are present [13] - for example, if multiple GATA1 motifs exist in an input sequence, but only one GATA1 motif is needed for the output to be positive, perturbing any single GATA1 motif may not cause a substantial change in the output. Such nonlinear relationships can be learned by the gkmrbf or wgkmrbf kernels (**Sec. 2.3**), both of which frequently produced improvements in performance relative to the gkm kernel [8].

SHAP [12] is a general-purpose interpretation method that addresses the saturation issue of ISM by sampling several combinations of input perturbations and looking at the change in the output in each case. Although SHAP has good theoretical guarantees, it is far less computationally efficient than ISM as the number of sampled perturbations needed to get accurate importance estimates can be quite large. For example, the authors of SHAP used 50, 000 sampled perturbations to obtain stable estimates of feature importance on a single example from the MNIST digit recognition dataset (each example is a 28 *×* 28 image).

## 4 Methods

### 4.1 Integrated gradients for gapped *k*-mer SVMs

In this section, we describe the gkmexplain method. Gkmexplain can be motivated in terms of the Integrated Gradients attribution method [15]. For any output function *F* (*X*), where *X* is a vector in ℝ*n*, and a path function *π* : [0, 1] *→* ℝ*n*, the *Path Integrated Gradients* [15] importance on the *i*th feature *X_i_* is defined as:

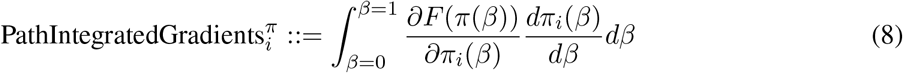

Equation 8 requires the output function *F* to be differentiable. Unfortunately, this is not the case for SVMs that use string kernels because DNA sequence is discrete. We solve this issue by introducing a collection of continuous variables 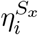 that can be interpreted as representing the “intensity” of the base at position *i* of sequence *S_x_*. Reformulating the SVM objective to contain these artificial intensity variables, we can apply the method of Integrated Gradients. A derivation is provided in **Appendix A.1**. The resulting formula for the importance of base *i* in sequence *S_x_* in a wgkm-SVM is:

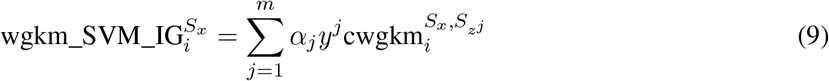

Where 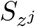, *α_j_* and *y^j^* are, respectively, the sequence, weight and label of support vector *j*, and:

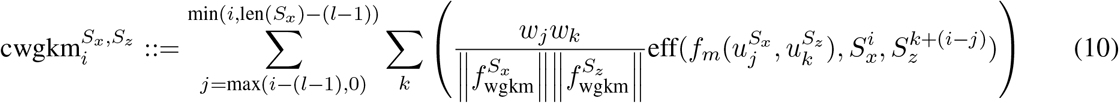

Where 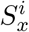 is the identity of the base at position *i* in sequence *S_x_* and

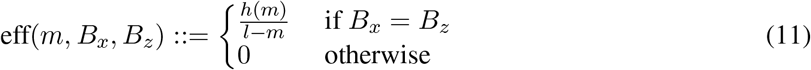

The formula for the importance of base *i* in position *S_x_* for the wkgmrbf kernel is:

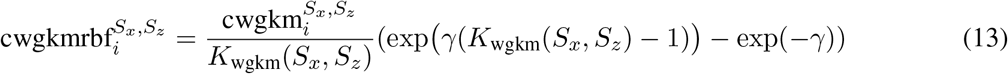

Where

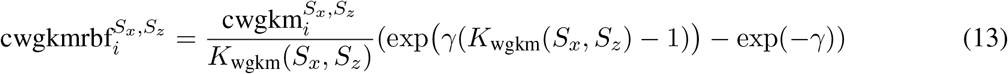

In terms of implementation, the gkmexplain importance scores can be computed efficiently by modifying the *k*-mer tree depth-first search that was originally used to compute the output of the gkm-SVM (see our implementation at https://github.com/kundajelab/lsgkm).

#### 4.1.1 Intuitive motivation

Although **Eqns. 9 & 12** can be justified using the method of Integrated Gradients, it can be helpful to have an intuitive understanding of the formulas. Recall from **Eqn. 3** that 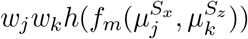 is the contribution of the *l*-mer 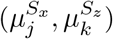 to the dot product 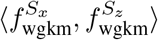. Let us denote 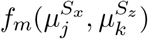as *m*, and let us consider a position *i* in sequence *S_x_* that overlaps 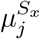. Let us denote the base at position *i* as 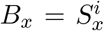, and the corresponding base within *l*-mer 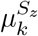 as 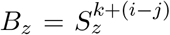 (“corresponding” means that the offset of *B_x_* relative to the start of 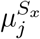 is the same as the offset of *B_z_* relative to the start of 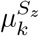). If we evenly distribute *w_i_w_j_h*(*m*) among the (*l − m*) matching positions between 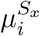 and 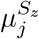, then position *i* would inherit an importance of 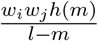 if *B_x_* = *B_z_*, and would inherit an importance of 0 if *B_x_≠B_z_*. In other words, using the definition of eff (*m, B_x_, B_z_*) in **Eqn. 11**, we can write the importance inherited by position *i* from the *l*-mer pair 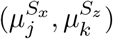 as 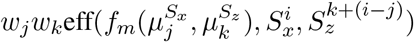 If we sum this quantity over all possible *l*-mer pairs where 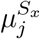 overlaps position *i*, and normalize by the total wgkm feature vector lengths (as is done in the wgkm kernel), we arrive at the quantity cwgkm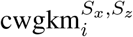 in **Eqn. 10**, which we can view as the contribution of base *i* in sequence *S_x_* to the kernel output *K*_wgkm_(*S_x_, S_z_*). The final importance of base *i* in **Eqn. 9** can then be seen as a weighted sum of the contributions of *i* to the kernel outputs 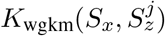, where the weights *α_j_ y^j^* are the same as the SVM weights.

As intuition for **Eqn. 12** (pertaining to the wgkmrbf kernel), note that exp(*−γ*) is the output of the *K*_wgkmrbf_(*S_x_, S_z_*) kernel when 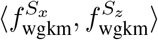 is 0. If we consider the “baseline” input *S_x_* to be one that has a wgkm dot product of 0 with all the support vectors, then 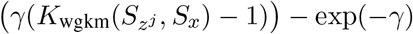 is the “difference from baseline” of *K*_wgkmrbf_(*S_x_, S_z_*). Earlier, we said that 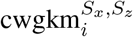 can be thought of as the contribution of base *i* in sequence *S_x_* to the kernel output *K*_wgkm_(*S_x_, S_z_*). The quantity 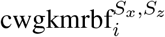 (**Eqn. 13**) can thus be viewed as attributing the “difference from baseline” of *K*_wgkmrbf_(*S_x_, S_z_*) to the positions *i* in proportion to the contribution of base *i* to *K*_wgkm_(*S_x_, S_z_*). The final importance of base *i* in **Eqn. 12** can then be seen as a weighted sum of the contributions of *i* to the “difference from baseline" of the kernel outputs 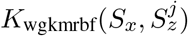, where the weights are *α_j_ y^j^*.

### 4.2 Mutation Impact Scores

The typical approach to estimating the impact of individual mutations is in-silico mutagenesis (ISM). In ISM, a mutation is introduced in the sequence and the change in the predicted output is computed. However, as illustrated in **Fig. 1**, ISM can overlook motifs if the response of the model has saturated in the presence of the motif (as can happen when the gkmrbf kernel is used). For these reasons, it can be beneficial to estimate the effects of mutations using a different approach. Building on **Sec. 4.1**, we introduce the term 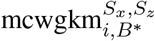 to represent the estimated impact on the kernel output *K*_wgkm_(*S_x_, S_z_*) of changing position *i* in sequence *S_x_* to base *B.^*^* We have:

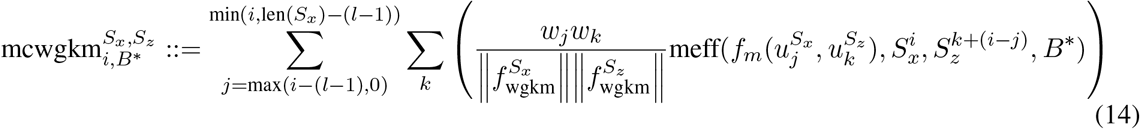

where

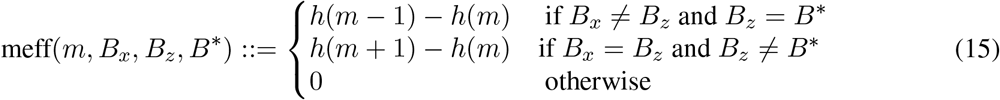

**Figure 1:**
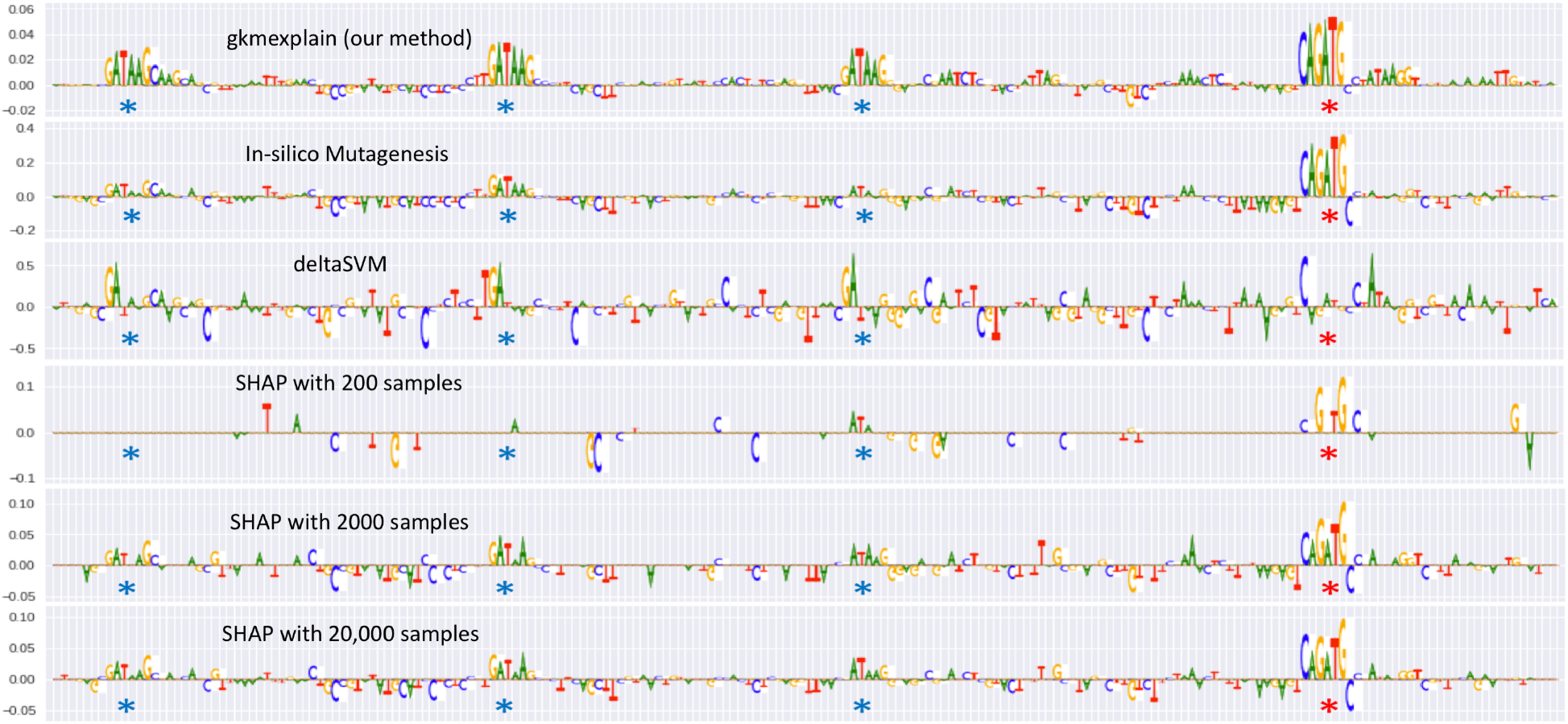
Comparison of gkmexplain, ISM, deltaSVM and SHAP on an individual sequence. An SVM with a gkmrbf kernel with *l* = 6, *k* = 5 and *d* = 1 was used to distinguish sequences containing both TAL1 and GATA1 motifs from sequences containing only one kind of motif or neither kind of motif. The locations of embedded GATA1 motifs are indicated by blue stars, and the location of the embedded TAL1 motif is indicated by a red star. For SHAP, a background of 20 shuffled versions of the original sequence was used. The relatively poor performance of ISM and deltaSVM is due to the nonlinear nature of the RBF kernel.

By analogy to **Eqns. 9 & 12**, we then define the mutation impact scores for base *B^*^* at position *i* in sequence *S_x_* as follows:

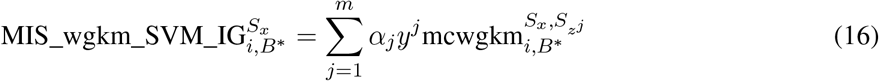

and

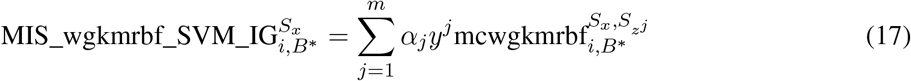

where

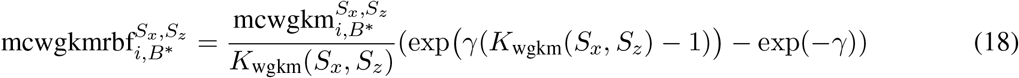

Once again, these quantities can be efficiently computed by modifying the *k*-mer tree depth-first search originally used to compute the output of the gkm-SVM (see our implementation at https://github.com/kundajelab/lsgkm). While the original implementation only performs recursion on *l*-mer pairs for which no more than *d* mismatches have been encountered so far, we perform recursion on *l*-mer pairs for which we may have encountered *d* + 1 mismatches (because a mutation can flip a mismatching position to a match).

## 5 Results

### 5.1 Simulated regulatory DNA sequences

To evaluate the performance of different explanation methods, we used the simulated genomics dataset from [13]. Briefly, 8000 200bp-long sequences were generated by randomly sampling the letters ACGT with background probabilities of 0.3, 0.2, 0.2 and 0.3 respectively. 0-3 instances of the TAL1_known1 and GATA1_disc1 motifs [7] were then embedded into non-overlapping positions in each sequence. 25% of sequences contained both embedded TAL1 motifs and embedded GATA1 motifs and were labeled +1.

The remaining sequences contained either just embedded GATA1 motifs, just embedded TAL1 motifs, or neither embedded TAL1 nor embedded GATA1 motif, and were labeled -1. 10% of sequences were reserved for a testing set, while the remaining were used for training. A Support Vector Machine with a gkmrbf kernel and parameters *l* = 6, *k* = 5 and *d* = 1 was trained to distinguish the positive set from the negative set. It attained 90% auROC. A notebook demonstrating model training and interpretation is at https://github.com/kundajelab/gkmexplain/blob/master/lsgkmexplain_TALGATA.ipynb.

**Fig. 1** illustrates the behavior of different algorithms on a sequence containing 3 GATA1 motifs and 1 TAL1 motif. For deltaSVM and ISM, the importance of a position was computed as the negative of the mean impact of all 3 possible mutations at that position (positions that produce negative scores when mutated will therefore receive positive importance). The gkmexplain method successfully highlights all GATA1 and TAL1 motifs present in the sequence. DeltaSVM performs very poorly, likely due to the nonlinear nature of the gkmrbf decision function (the nonlinearity is needed to learn the logic that both GATA1 and TAL1 must be present in the sequence for the output to be positive). ISM fails to clearly highlight some individual GATA1 motifs in this sequence, likely because the presence of multiple GATA1 motifs has a saturating effect on the nonlinear decision function. To confirm this was not an isolated example of the failure of ISM, we compared the ability of gkmexplain and ISM to identify the embedded motifs across 2000 examples, and found that gkmexplain does indeed perform better (**Fig. 3 & 4**). SHAP shows promise at highlighting the relevant motifs, but only when many perturbation samples are used. Unfortunately, when a large number of perturbation samples are needed, SHAP has a very slow runtime compared to the other methods (**Fig. 2**).

**Figure 2:**
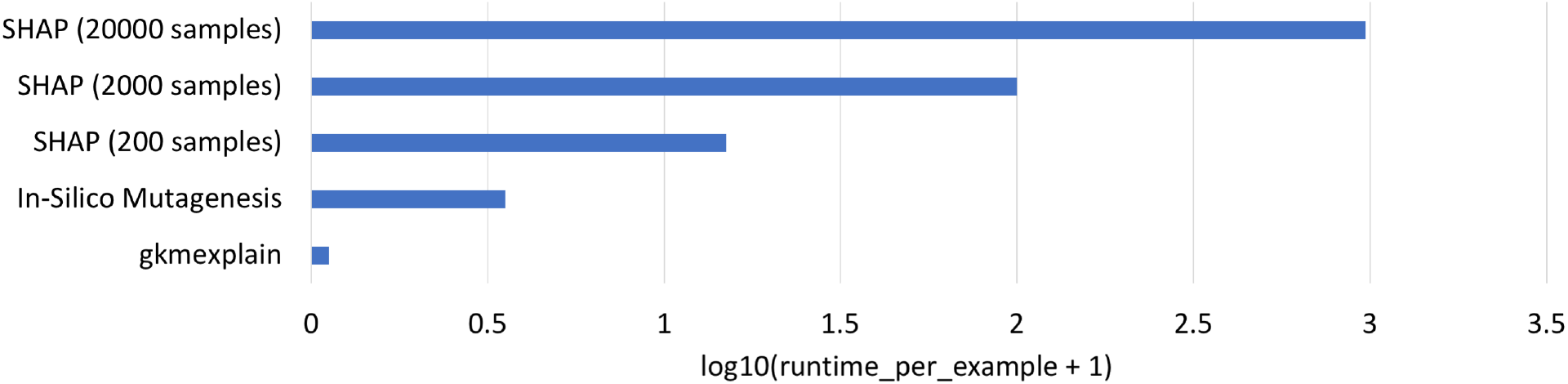
Time taken to compute importance on a single sequence for various algorithms (log scale). Evaluation was done on the model and data described in 5.1. For SHAP, a background of 20 shuffled versions of the original sequence was used (the runtime of SHAP scales linearly in the product of the size of the background and the number of samples).

**Figure 3:**
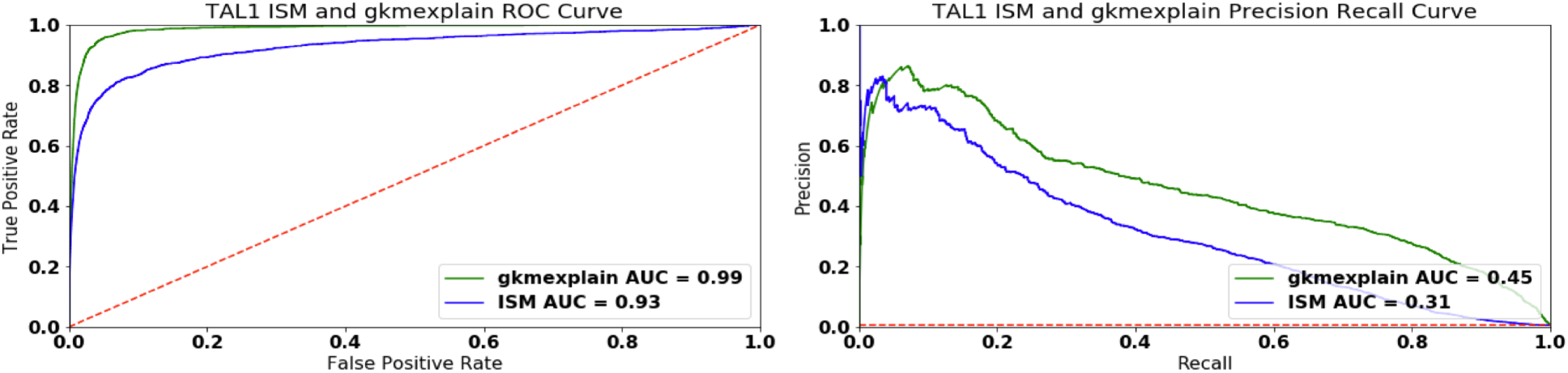
gkmexplain outperforms ISM at identifying TAL1 motifs. A gkmrbf SVM was trained as described in **Sec. 5.1**, but with a random heldout set of size 2000 rather than 800. The 2000 heldout examples were scanned using 16bp windows (the length of the TAL1_known1 motif). Windows containing a complete embedded TAL1_known1 motif were labeled positive. Windows containing no portion of any embedded motif were labeled negative. All other windows were excluded from analysis. Windows were ranked according to the total importance score produced by the importance scoring method in question, and AuROC and AuPRC were computed. The gkmexplain method outperforms ISM on both metrics. The dashed red line shows the performance of a random classifier.

**Figure 4:**
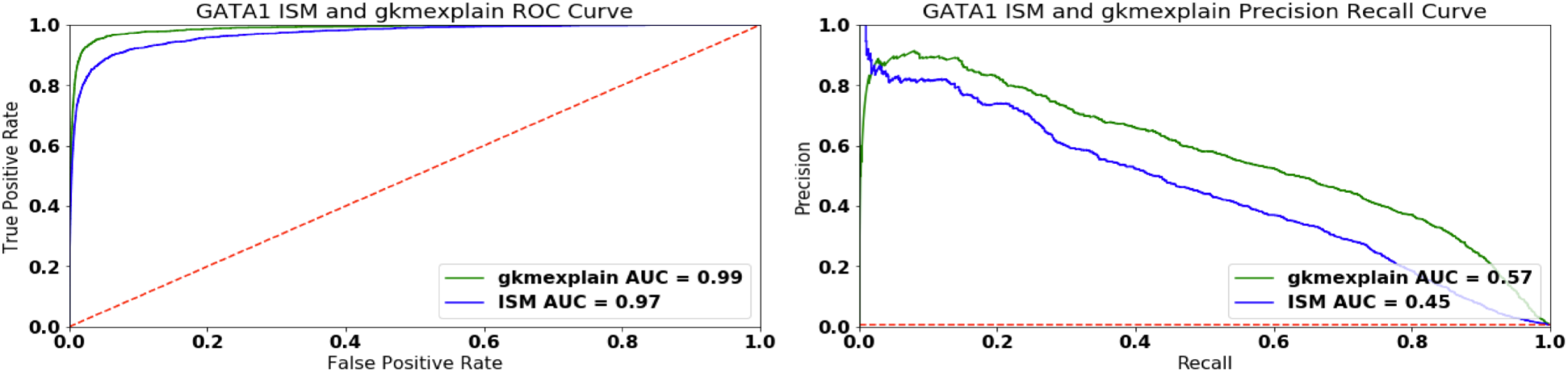
gkmexplain outperforms ISM at identifying GATA1 motifs. Analogous to **Fig. 3**, but using 10bp windows (the length of the GATA1 motif) rather than 16bp windows.

As further confirmation that gkmexplain was able to detect the embedded TAL1 and GATA1 motifs, we supplied gkmexplain-derived importance score profiles across all sequences in the positive set a new motif discovery method called TF-MoDISco [14]. Briefly, TF-MoDISco identifies subsequences (termed “seqlets”) of high importance in all the input sequences, builds an affinity matrix between seqlets using a cross-correlation-like metric, clusters the affinity matrix, and then aggregates aligned seqlets in each cluster to form consolidated motifs. TF-MoDISco accepts both importance score profiles as well as “hypothetical” importance score profiles of multiple sequences. Hypothetical importance scores can be intuitively thought of as revealing how the classifier might respond to seeing alternative bases at any given position in a sequence. For gkmexplain, we derived these hypothetical scores using the method described in **Appendix A.2**. We also normalized the scores as described in **Appendix A.2.1**. The resulting motifs are shown in **Fig. 5**. We find that TF-MoDISco is able to learn motifs that closely match the true embedded motifs.

**Figure 5:**
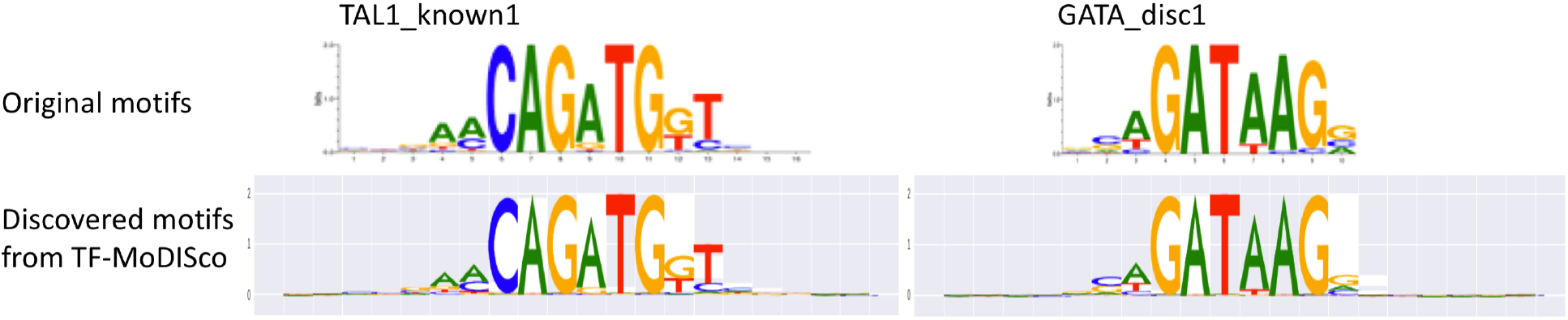
Motifs extracted by running TF-MoDISco on gkmexplain importance scores successfully recovers ground-truth simulated motifs. Letter heights are proportional to the information content of the probabilities across the different bases at that position.

### 5.2 NFE2 *in vivo* Transcription Factor binding models

We also studied the performance of gkmexplain on a gkmSVM model trained on ChIP-seq data for the NFE2 transcription factor in Gm12878. The model and example sequences were directly obtained from the published lsgkm repository [10]. The model was trained using a wgkm kernel with *l* = 10, *k* = 6 and *d* = 3 to discriminate 644 variable-length DNA sequences that overlap NFE2 ChIP-seq peaks in GM12878 from an equal number of genomic background sequences matched for GC-content and repeat-fraction. Gkmexplain importance scores were then computed on the 644 positively-labeled training sequences and 69 positively-labeled test sequences that shipped with the repository. Scores on a few example sequences are shown in **Fig. 6**. The results of TF-MoDISco are in **Fig. 7**. TF-MoDISco successfully recovers the top two motifs for NFE2 identified using traditional motif discovery methods [1, 16], as well as some evidence of a motif dimer. A notebook reproducing the results is at https://github.com/kundajelab/gkmexplain/blob/master/lsgkmexplain_NFE2.ipynb.

**Figure 6:**
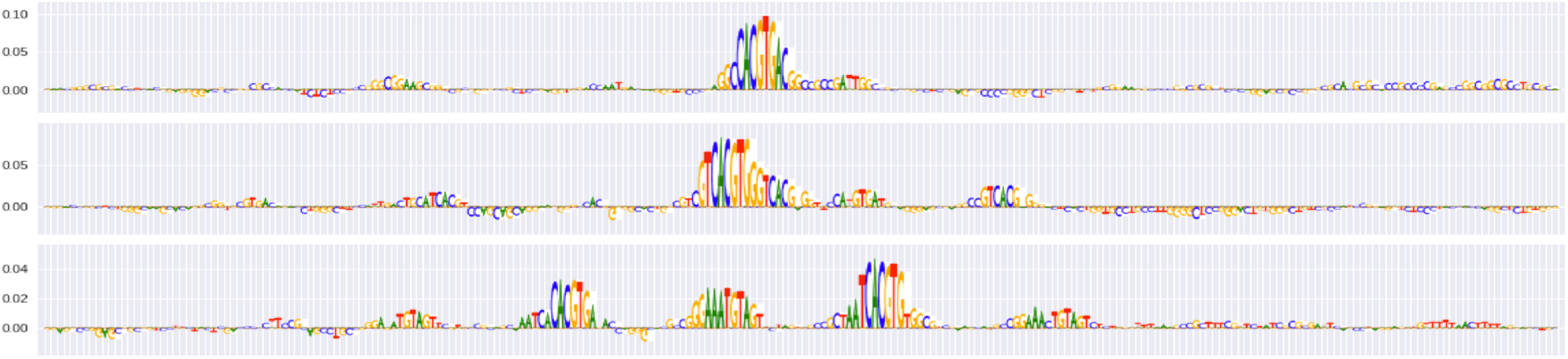
Gkmexplain importance scores on example NFE2-bound sequences.

**Figure 7:**
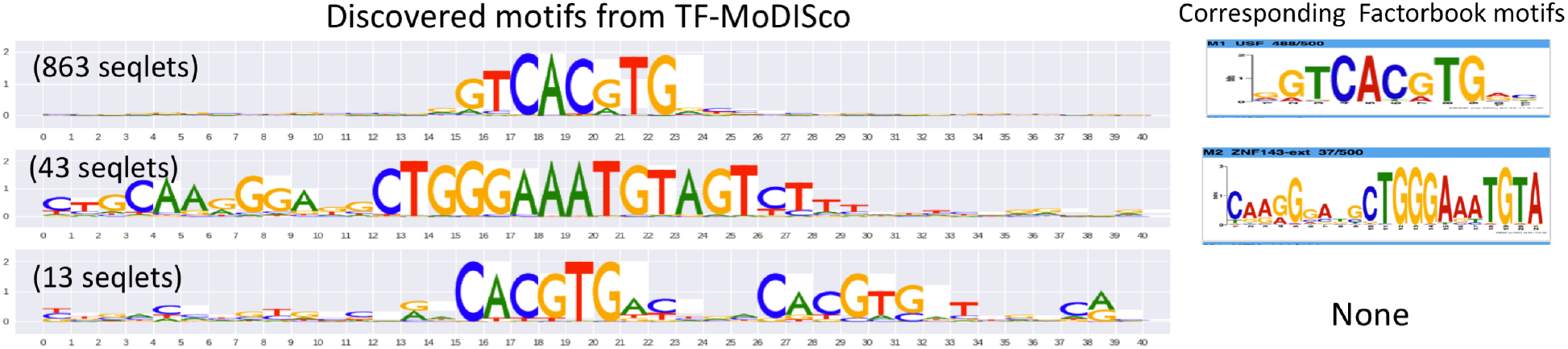
NFE2 motifs derived using TF-MoDISco. Letter heights are proportional to the information content of the probabilities across the different bases at that position.“Seqlets” are subsequences of high importance that are used by TF-MoDISco to create motifs [14]. The number of seqlets contained within each TF-MoDISco motif is indicated; motifs with a larger number of seqlets are more reliable.

### 5.3 Predicting regulatory genetic variants affecting DNase hypersensitivity

We trained models using the lsgkm package [8] on a DNase-seq dataset in the GM12878 lymphoblastoid cell-line. The training dataset is publicly downloadable from the deltaSVM website [3]. It consists of a single positive set containing 22,384 300bp sequences that overlapped DNase hypersensitive peaks, and five independently-generated negative sets that each matched the size, length distribution, GC-content and repeat-fraction of the positive set. One gkm-SVM and one gkmrbf-SVM were trained for each choice of negative set. For the gkm-SVM and gkmrbf-SVM, we used the parameter settings *l* = 10, *k* = 6 and *d* = 3, consistent with the deltaSVM paper. For gkmrbf-SVM, we further set the regularization parameter *c* to 10 and the gamma value *g* to 2, as suggested by the lsgkm documentation [10]. Remaining values were left to the lsgkm defaults.

To assess whether gkmexplain could be used to quantify the functional impact of regulatory genetic variants, we used the same benchmarking dataset of DNase I–sensitivity quantitative trait loci (dsQTLs) in lymphomablastoid cell lines (LCLs) that was used in the deltaSVM paper [9, 5], consisting of 579 dsQTL SNPs and 28,950 control SNPs. The dsQTL SNPs were each located within their associated 100bp DNase hypersensitive peak and had an association *p*-value below 10^*−*5^, while the control SNPs each had minor allele frequency above 5% and were randomly sampled from the top 5% of DNase hypersensitive sites.

We used 4 methods to score dsQTLs and control SNPs: deltaSVM applied to the gkm-SVM, deltaSVM applied to the gkmrbf-SVM, in-silico mutagenesis (ISM) applied to the gkmrbf-SVM, and gkmexplain mutation impact scores (**Sec. 4.2**) applied to the gkmrbf-SVM. For ISM and gkmexplain, a window of 51bp centered on the SNP was used as context. Results are shown in **Fig. 8**. Across the models trained on the 5 independent negative sequence sets, gkmexplain consistently produces the best auPRC (binomial p-value = 0.5^5^ = 0.03125). Interestingly, we found that deltaSVM applied to the gkmrbf-SVM consistently produced better auRPC than deltaSVM applied to the gkm-SVM, even though Lee found that deltaSVM did not produce improvements when used with the gkmrbf kernel [8], possibly due to differences in the dataset and parameter settings. Results for auROC are in **Appendix A.3**.

**Figure 8:**
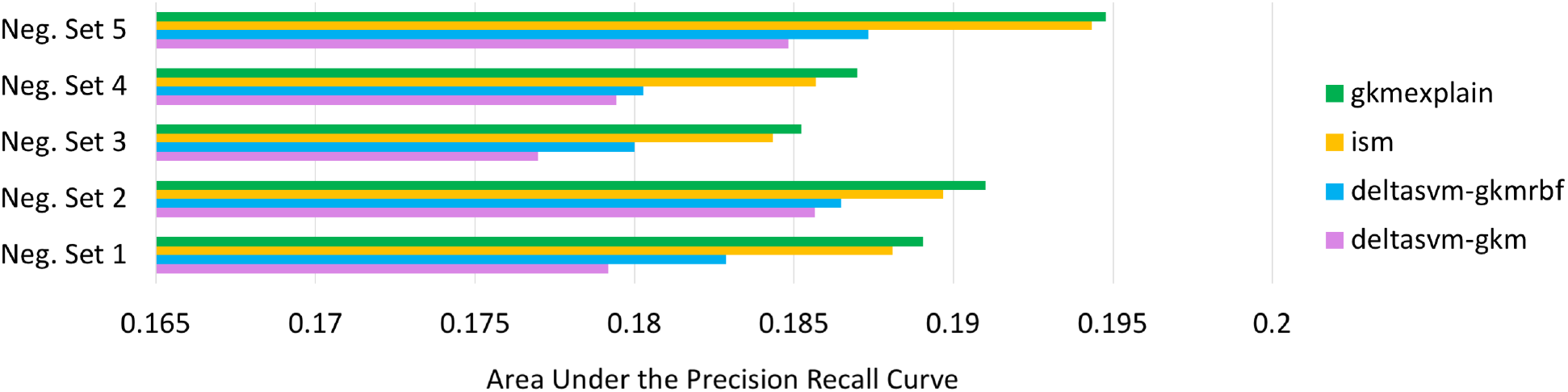
Gkmexplain Mutation Impact Scores Outperform deltaSVM and ISM at identifying dsQTLs. For each choice of negative set provided in the deltaSVM paper, we trained a gkm-SVM and gkmrbf-SVM. LCL dsQTLs and control SNPs were then scores using four methods: deltaSVM on the gkm-SVM, deltaSVM on the gkmrbf-SVM, ISM on the gkmrbf-SVM and gkmexplain Mutation Impact Scores (**Sec. 4.2**) on the gkmrbf-SVM. For ISM and gkmexplain, a 51bp window centered around the SNP was used as context. The gkmexplain mutation impact scores consistently produce the best auPRC across all 5 choices of the negative set (binomial p-value = 0.5^5^ = 0.03125).

## 6 Conclusion

We have presented gkmexplain, an algorithm based on Integrated Gradients that can be used to efficiently explain the predictions of a Support Vector Machine trained with various gapped *k*-mer string kernels. On simulated data, we showed that our method outperforms ISM and deltaSVM when used with a nonlinear gapper *k*-mer kernel such as the gkmrbf kernel (**Fig. 1, 3 & 4**), while being far more computationally efficient than ISM or SHAP (**Fig. 2**). We further demonstrated that importance scores derived through gkmexplain can be supplied to TF-MoDISco [14] to perform motif discovery (**Fig. 5 & 7**). On a DNAse-I hypersensitivity QTL dataset derived from lymphomablastoid cell lines, we found that gkmexplain Mutation Impact Scores consistently produced better auPRC than ISM or deltaSVM (**Fig. 8**). Finally, we note that the idea of using Integrated Gradients to interpret SVMs is not limited to genomics; it can be applied to other kinds of data, as illustrated in **Appendix A.4**.

## A Appendices

### A.1 Derivation of Equations 9 and 12

This section derives the formulas present in **Sec. 4.1** using the method of Integrated Gradients.

Applying the definition of Path Integrated Gradients from **Eqn. 8** to the objective function for SVMs in **Eqn. 1** we get:

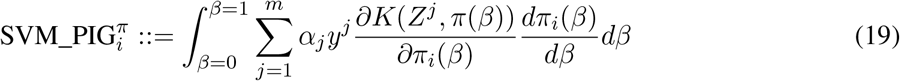

As mentioned earlier, the primary hurdle when applying Integrated Gradients to support vector machine string kernels is that the formula for Integrated Gradients requires the kernel function *K* to be differentiable. Because DNA sequence is discrete, most string kernels are not differnentiable. To circumvent this issue, we introduce vector of variables 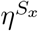 where each element 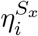 can be interpreted as the “intensity” of the base at position *i* of sequence *S_x_*. Let 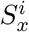 denote the identity of base *i* in sequence *S_x_*, let *l* be the length of the *l*-mers, and let 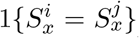 be an indicator function that takes the value of 1 if 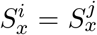 and 0 otherwise. We rewrite the wgkm kernel between an input sequence *S_x_* and a support vector sequence *S_z_* as a function of 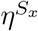 as follows:

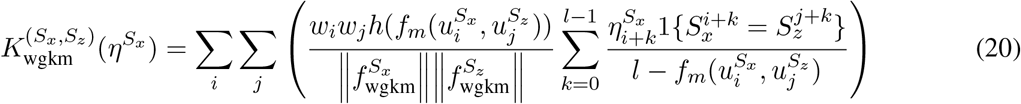

When 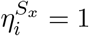 for all *i*, 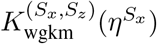 is equal to *K*_wgkm_(*S_x_*, *S_z_*).

Recall that 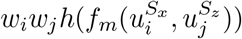 is the contribution of the *l*-mer pair 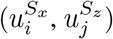 to 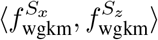. In the expression above, 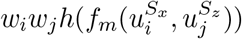 is scaled according to the total intensity of all the 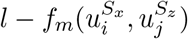 matching bases between 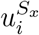and 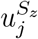.

Let len(*S_x_*) be a function that returns the length of the sequence *S_x_*. Because **Eqn. 20** is differentiable, we can compute the partial derivative 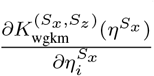 as:

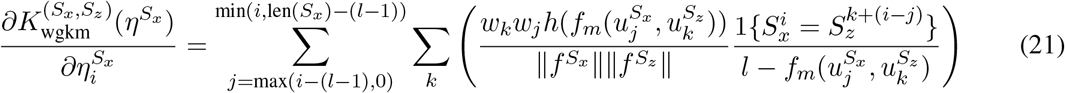

Intuitively, the expression looks at each *l*-mer 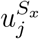 in *S_x_* that overlaps position *i*, compares it to every *l*-mer 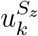 in *S_z_*, and increments the gradient proportionally to 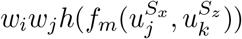 only if the base at position (*i − j*) in 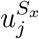 is a match to the corresponding base in 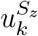 . Note that the number of *l*-mers in sequence *S_x_* is (len(*S_x_*) *−* (*l −* 1))

Rewriting the formula for Path Integrated Gradients in **Eqn. 19** using the differentiable reparameterization in **Eqn. 20**, we get:

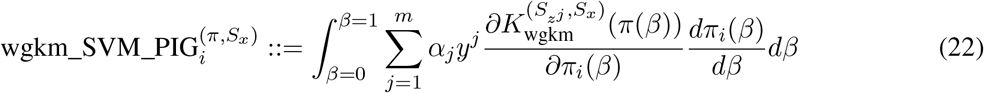

If we now consider the path *π_i_*(*β*) = *β* (i.e. if we linearly scale the intensities 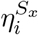 uniformly between 0 and 1), we would have 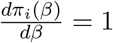 . This choice of path is the standard “Integrated Gradients” path. Because the partial derivative in **Eqn. 21** is constant with respect to 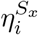, simplifying the expression above results in the formula for the importance of individual bases in a wkgm SVM that was presented earlier in **Eqn. 9**, reproduced below for convenience:

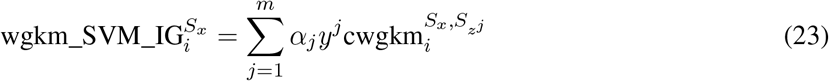

Where 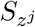, *α_j_* and *y^j^* are, respectively, the sequence, weight and label of support vector *j*, and:

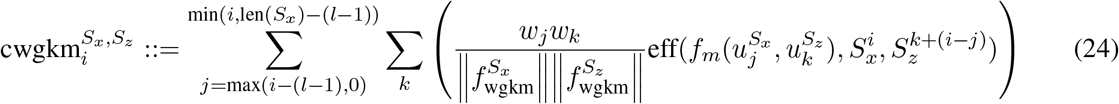

Where 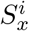 is the identity of the base at position *i* in sequence *S_x_* and

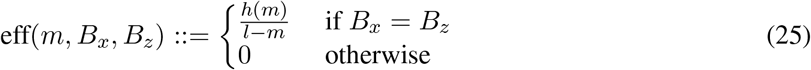

For the case of the wgkmrbf kernel (**Eqn. 7**), we once again reparameterize the kernel in terms of 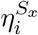 as follows:

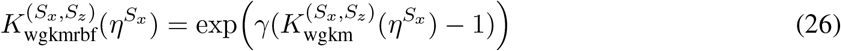

By the chain rule, we get:

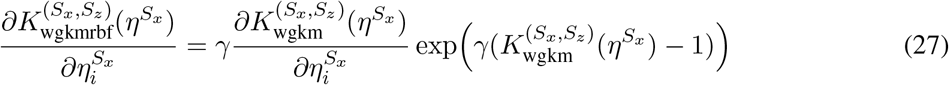

As before, we consider the path *π_i_*(*β*) = *β* (i.e. we linearly scale the intensities 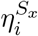 uniformly between 0 and 1). From **Eqn. 20**, we note that 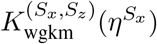is linear in the values of 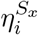 . Thus, we can write:

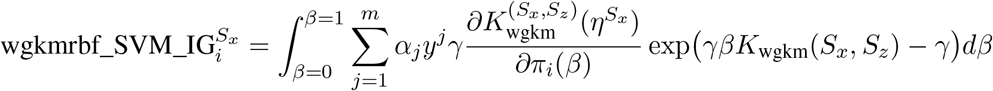

Recall that 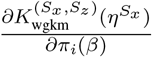 is constant with respect to *π_i_*(*β*). After performing the integral with respect to *β*, we obtain **Eqn. 12**, reproduced below for convenience:

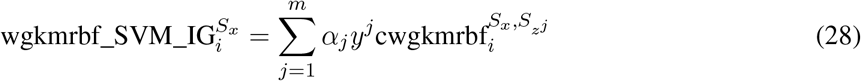

Where

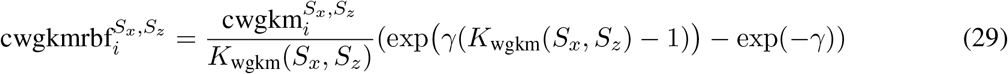

### A.2 Hypothetical importance scores

For motif discovery with TF-MoDISCo [14], it is useful to have *hypothetical* importance scores in addition to the true importance scores. The hypothetical importance score of base *B* at position *i* estimates the preference of the classifier for seeing base *B* at position *i* instead of the base that is actually present at position *i*. If base *B* is the same as the base that is actually present in the sequence at position *i*, the hypothetical importance score is defined to be the same as the actual importance score. As an example, suppose a particular TF has high affinity to the sequences GATAAT and GATTAT and low affinity to the sequences GATCAT and GATGAT. If the sequence GATAAT is presented to a classifier trained to predict binding sites of the TF, the hypothetical importance assigned to base T in the 4th position would be positive, as would be the (actual) importance assigned to base A in the 4th position. By contrast, the hypothetical importance scores assigned to C and G in the 4th position would be negative. To compute hypothetical importance scores, we first define the quantity 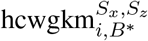 (analogous to 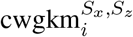 in **Eqn. 10**) as follows:

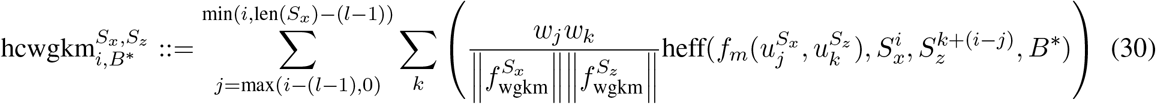

where

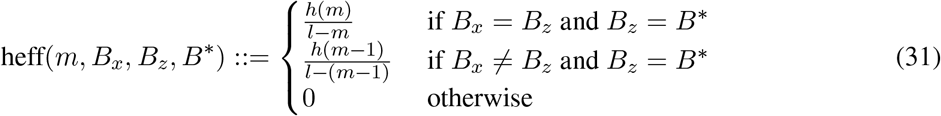

By analogy to equations 9 and 12, we then define the hypothetical contribution scores for base ^*B\**^ at position *i* in sequence *S_x_* as follows:

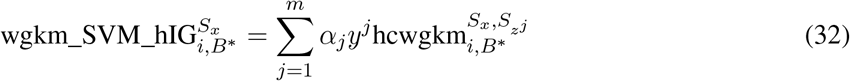

and

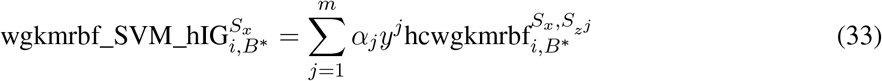

where

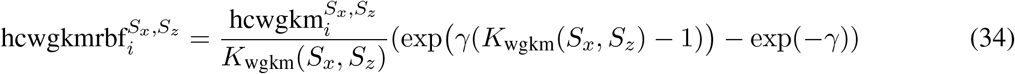

The hypothetical importance scores are useful for motif discovery with TF-MoDISco because different instances of a motif that have variations in their underlying sequence may nonetheless have similar hypothetical importance scores because the hypothetical importance scores impute the importance on all possible bases that could be present (and not just the bases that happen to be present in the specific instance of the motif). If the goal is to infer the impact of individual mutations rather than to perform motif discovery, the Mutation Impact Scores (described in **Sec 4.2**) would be more appropriate.

Hypothetical importance scores can be efficiently computed by modifying the *k*-mer tree depth-first search originally used to compute the output of the gkm-SVM (see our implementation at https://github.com/kundajelab/lsgkm). While the original implementation only performs recursion on *l*-mer pairs for which no more than *d* mismatches have been encountered so far, in order to get the most accurate hypothetical importance scores we should perform recursion on *l*-mer pairs for which we may have encountered *d* + 1 mismatches because a mutation can flip a mismatching position to a match. However, the additional layer of recursion can increase runtime substantially. In practice, we found that hypothetical importance scores derived by only considering recursions on *l*-mer pairs with up to *d* mismatches work well, and that is what we used in this paper.

#### A.2.1 Normalization of scores for TF-MoDISco

Empirically, we found that the following normalization of the importance scores produces improved results with TF-MoDISco. Let *f_h_*(*S_x_, i, B^*^*) be a function that returns the hypothetical importance of base *B^*^* at position *i* in sequence *S_x_*, and let 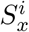 be the base at position *i* in sequence *S_x_*. To normalize the hypothetical importance scores at position *i*, we divide by the sum of all hypothetical scores at position *i* with the same sign as 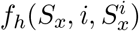. The rationale is that if a different base at position *i* could produce a score of higher magnitude than 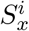, then 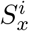is relatively less important. Let 1{*x >* 0} be an indicator function that returns 1 if *x* is positive and 0 otherwise. Formally, the normalized hypothetical importance for base *B^*^* at position *i* is defined as:

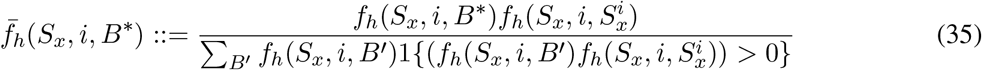

Similarly, we define the normalized importance score as:

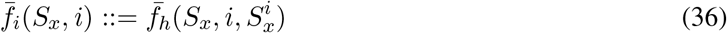

We find that the normalized importance scores appear to be less noisy relative to the unnormalized importance scores, as illustrated in **Fig. A1**.

**Figure A1:**
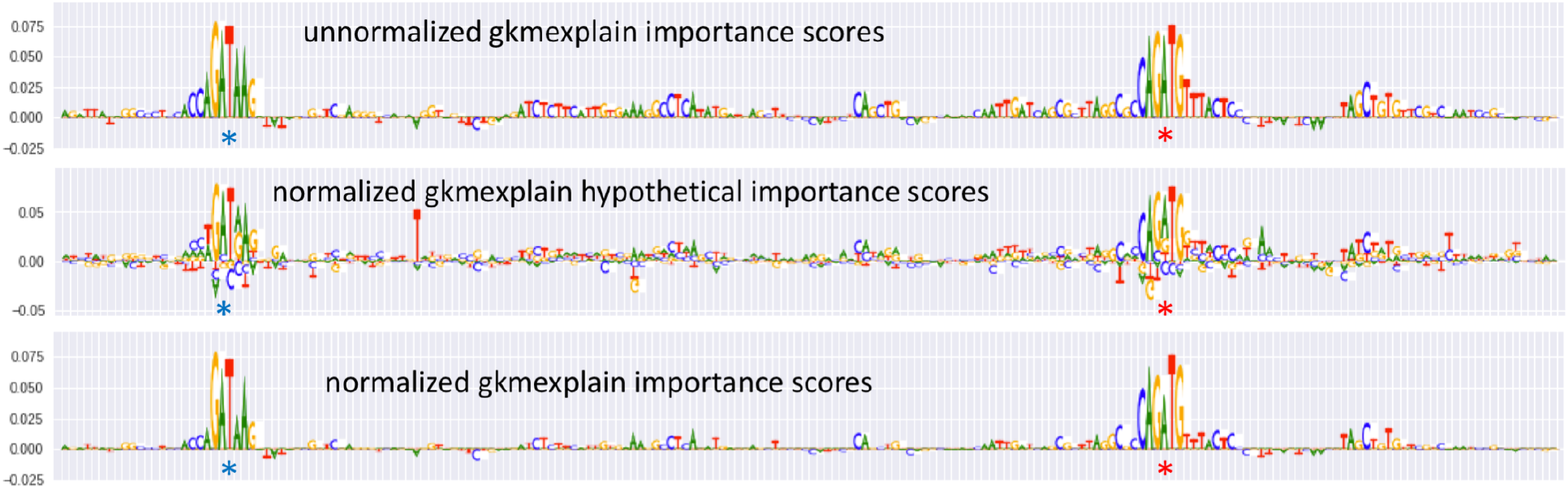
Normalization of gkmexplain importance scores for TF-MoDISco. A gkmrbf SVM was trained as described in **Sec. 5.1**. Shown are unnormalized gkmexplain importance scores (top row), normalized hypothetical importance scores (**Eqn. 35**), and normalized importance scores (**Eqn. 36**) on a single sequence. The blue star indicates the location of an embedded GATA1 motif (GATAAG), and the red star indicates the location of an embedded TAL1 motif (CAGATG). The normalized importance scores appear less noisy than the unnormalized importance scores.

### A.3 AuROC on dsQTL data

Performance in terms of auROC on the dsQTL dataset is shown in **Fig. A2**. Although gkmexplain gives a weaker auROC than ISM in 4 out of 5 cases, the difference is not statistically significant (possibly owing to the small number of samples; binomial p-value = 0.1875). We note that when negatives greatly outnumber positives, auPRC is generally considered a more useful performance metric than auROC. We also note that gkmexplain consistently outperforms deltaSVM.

**Figure A2:**
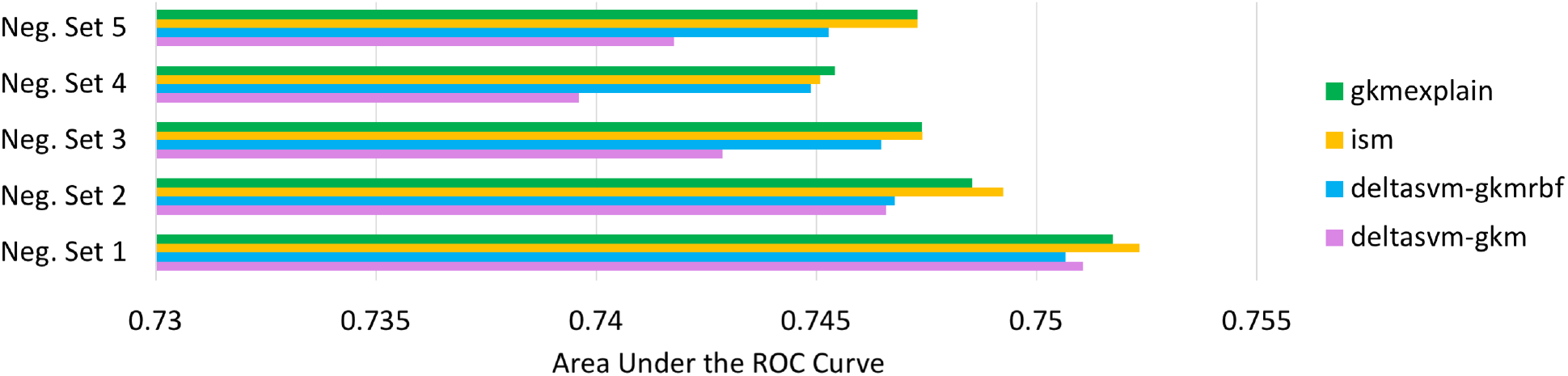
AuROC for dsQTL identification. Analogous to **Fig. 8**, but showing auROC instead of auPRC. The gkmexplain and ISM methods consistently outperform deltaSVM.

### A.4 Integrated Gradients for interpreting other non-linear SVM models

To explore the general applicability of Integrated Gradients to SVM interpretation beyond genomics, we used the MADELON dataset [2]. This dataset has a training set of 2000 examples and a validation set of 600 examples, all with 500 features, of which 20 features are informative and the remaining are “distractor" features with no predictive power. Because the Gaussian SVM had a strong tendency to overfit to the full dataset, we created 10 reduced feature sets, each with the same set of examples in the training and validation sets, but with 20 likely informative features (identified based on the challenge submission results) and a different set of 20 distractor features in each set. Each of these 10 sets served as a distinct data point on which to evaluate the accuracy of the importance scoring algorithm in question (Integrated Gradients or SHAP). Each feature in each set was normalized by subtracting the average feature value and dividing by the standard deviation. We trained Gaussian kernel SVM on each of the datasets and achieved approximately 0.8 accuracy on the validation set. These classifiers were then provided to the importance scoring algorithms.

When applying Integrated Gradients, we used a reference that was the average feature value of the negative labeled training points and approximated the integration of the partial derivatives using 10 linearly-spaced intermediate points between each test point and the reference. With SHAP, we experimented with 10 and 50 samples around the test points using the same reference as was used for IG. We judged the quality of each importance scoring algorithm by feeding the top 10 most important features reported by the algorithm to a Gaussian kernel SVM and measuring the SVM’s accuracy on the validation set. The results are shown in Figure A3. Integrated Gradients is significantly faster than SHAP and tends to be more accurate than when SHAP is used with a small number of perturbation samples. With a larger number of perturbation samples, SHAP is more accurate, though it is still slower. A notebook demonstrating how to use Integrated Gradients on a non-genomic dataset is at https://colab.research.google.com/drive/1LUGMIwOHLKDdK3deg7IIZ71kb2Nu1qv2.

**Figure A3:**
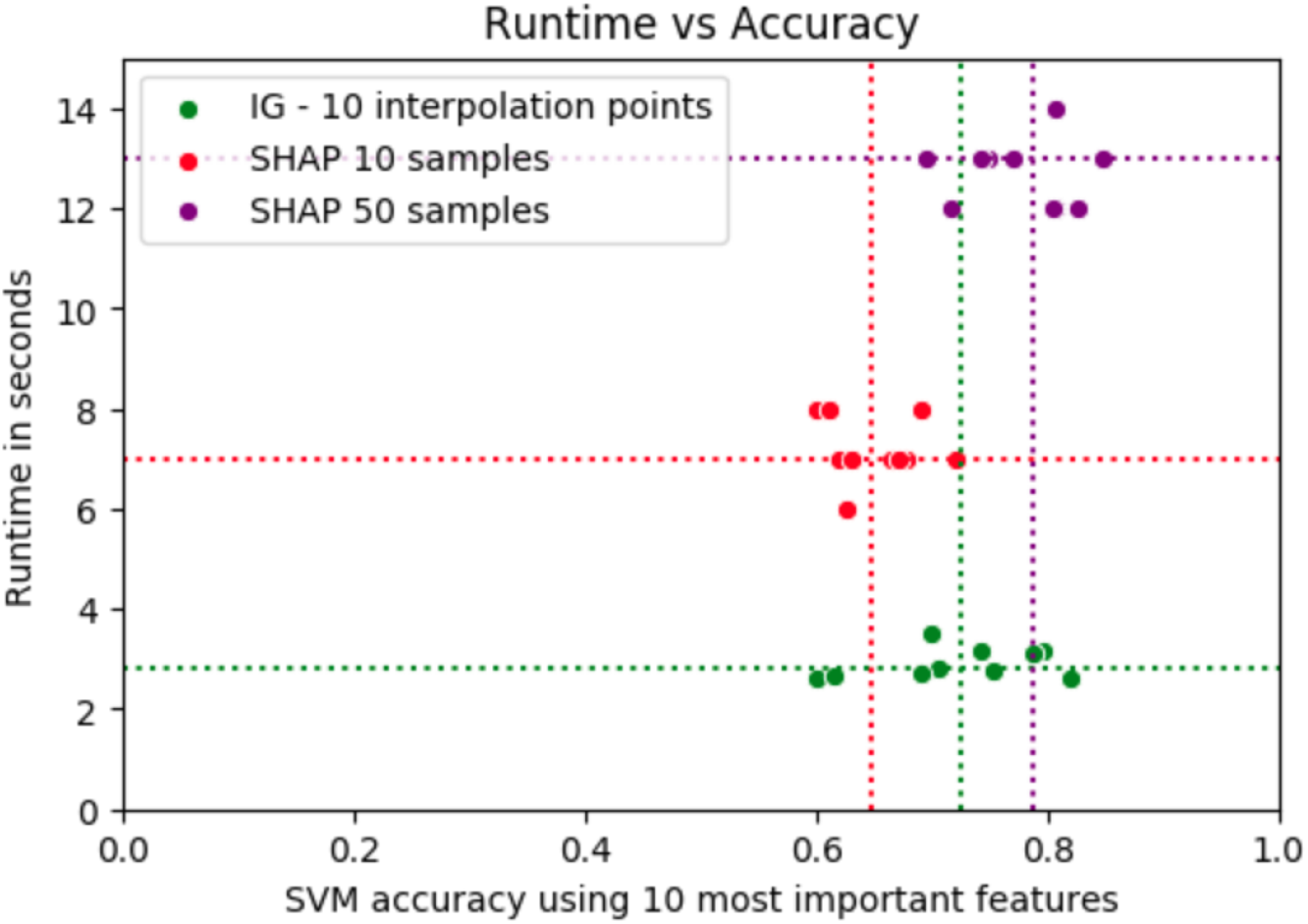
Comparison of Integrated Gradients with SHAP for Gaussian kernel SVM interpretation on the Madelon dataset. Dashed lines indicate means. IG appears to be pareto optimal in the sense that it can provide more accurate results faster than compared to SHAP used with a small number of perturbation samples.

